# Mitochondrial fission during mitophagy requires both inner and outer mitofissins

**DOI:** 10.1101/2025.04.13.648642

**Authors:** Kentaro Furukawa, Tatsuro Maruyama, Shun-ichi Yamashita, Keiichi Inoue, Tomoyuki Fukuda, Nobuo N. Noda, Tomotake Kanki

**Affiliations:** Department of Cellular Physiology, Graduate School of Medical Sciences, Kyushu University, Fukuoka 812-8582, Japan; Institute of Microbial Chemistry (BIKAKEN), Shinagawa-ku, Tokyo 141-0021, Japan; Department of Cellular Physiology, Niigata University Graduate School of Medical and Dental Sciences, Niigata 951-8510, Japan; Institute for Genetic Medicine, Hokkaido University, Sapporo, Hokkaido 060-0815, Japan

**Keywords:** Atg44, Dnm1, Mfi2, mitochondria, mitochondrial fission, mitofissin, mitophagy, yeast

## Abstract

Mitophagy maintains mitochondrial homeostasis through selective degradation of damaged or excess mitochondria. Recently, we identified mitofissin/Atg44, a mitochondrial intermembrane space-resident fission factor, which directly acts on lipid membranes and drives mitochondrial fission required for mitophagy in yeast. However, it remains unclear whether mitofissin is sufficient for mitophagy-associated mitochondrial fission and whether other factors act from outside the mitochondria. Here, we identify a mitochondrial outer membrane-resident mitofissin-like microprotein required for mitophagy, and we name it mitofissin 2/Mfi2 based on the following results. Overexpression of a C-terminally truncated form of Mfi2 induces mitochondrial fragmentation and partially restores mitophagy in *atg44*Δ cells. Mfi2 binds to lipid membranes and mediates membrane fission *in vitro*, demonstrating its intrinsic mitofissin activity. Genetic analyses reveal that Mfi2 and the dynamin-related protein Dnm1 independently facilitate mitochondrial fission during mitophagy. Thus, Atg44 and Mfi2, two mitofissins with distinct localizations, are required for mitophagy-associated mitochondrial fission.

## INTRODUCTION

Autophagy is a catabolic process that mediates vacuolar/lysosomal degradation and recycling of various cytoplasmic components, including organelles. Upon induction of autophagy, double-membranous structures called isolation membranes or phagophores emerge in the cytosol, expand, and engulf cytoplasmic proteins and organelles to form autophagosomes. The autophagosome fuses with vacuoles in yeast and plants or lysosomes in mammals, leading to degradation of its sequestered materials by vacuolar/lysosomal hydrolases.^1^ Mitophagy is a type of autophagy that selectively degrades damaged or excess mitochondria and contributes to mitochondrial homeostasis.^2–4^ Autophagy and mitophagy share several molecular processes, but the latter requires receptor-mediated recognition of particular mitochondrial regions as cargoes. Various mitochondrial outer membrane-resident receptors have been reported from yeast^5–7^ to mammals.^8–12^ The size of the mitochondrial cargo is also critical for mitophagy. Because mitochondria are typically larger than the autophagosome, mitochondria need to be fragmented to allow their engulfment within the autophagosome.^13–16^ However, the known mitochondrial fission factors, dynamin-related proteins Dnm1 in yeast and Drp1 in mammals, have been shown to be dispensable for mitophagy,^17–19^ suggesting the involvement of an alternative fission mechanism in this process.

Recently, we reported that the mitochondrial intermembrane space protein Atg44 (also known as Mdi1/Mco8) drives mitochondrial fission during mitophagy in the yeast *Saccharomyces cerevisiae*, and that Atg44 directly binds to lipid membranes and induces membrane fragility to facilitate membrane fission.^20^ Therefore, we named this protein “mitofissin” (*mito*chondrial *fis*sion prote*in*). In addition, Atg44 is required for the completion of Dnm1-mediated mitochondrial fission under homeostatic conditions,^21,22^ suggesting coordinated actions of these two fission factors from inside and outside the mitochondria, respectively. However, these findings do not fully explain why Dnm1 is dispensable for mitophagy. Moreover, it remains unclear whether Atg44 is sufficient for mitophagy-associated mitochondrial fission.

In this study, we identified a mitochondrial outer membrane-resident microprotein, Mco12, that is partially required for mitophagy. *In vivo* and *in vitro* analyses revealed that Mco12 possesses mitofissin activity, and we therefore renamed this protein “*m*ito*fi*ssin *2*” (Mfi2). Furthermore, we found that Mfi2 and Dnm1 facilitate mitochondrial fission in parallel from outside the mitochondria during mitophagy, providing an explanation for why disruption of Dnm1 alone does not prevent mitophagy. We propose that inner and outer mitofissins, Atg44 and Mfi2, together with Dnm1, coordinate mitophagy-associated mitochondrial fission from inside and outside the mitochondria.

## RESULTS

### Identification of Mco12/Mfi2 as an Atg44-like microprotein

Since mitofissin/Atg44 was one of the previously unexplored microproteins, we expected that other microproteins might also play a role in mitophagy. Therefore, we systematically examined mitophagy in yeast cells lacking mitochondrial micro- or small proteins identified in previous mitochondrial proteomic analyses^23^ (Table S1). In yeast, mitophagy can be induced either by nitrogen starvation (SD-N medium) or by continuous culture in a non-fermentable medium (YPL medium) to stationary phase, and can be monitored by assessing the vacuolar processing of chimeric mitochondrial proteins, such as Om45-GFP and Idh1-GFP, resulting in the release of free GFP.^5,24^ We found that among the mutant cells analyzed, *mco12*Δ (*mfi2*Δ) cells exhibited slightly decreased mitophagy upon nitrogen starvation compared with wild-type (WT) cells (Figure S1A). We confirmed this slight decrease by a time course assay (Figure 1A, *mfi2*Δ) and observed a similar decrease in mitophagy induced during the stationary phase (Figure 1B, *mfi2*Δ). Thus, we concluded that Mco12 is partially required for mitophagy. Next, we analyzed mitochondrial morphology by fluorescence microscopy in cells lacking mitochondrial micro- or small proteins in YPL medium. Unlike *atg44*Δ cells with enlarged mitochondria, the other mutant cells showed fragmented mitochondria similar to those in WT cells (Figure S1B).

**Figure 1.**
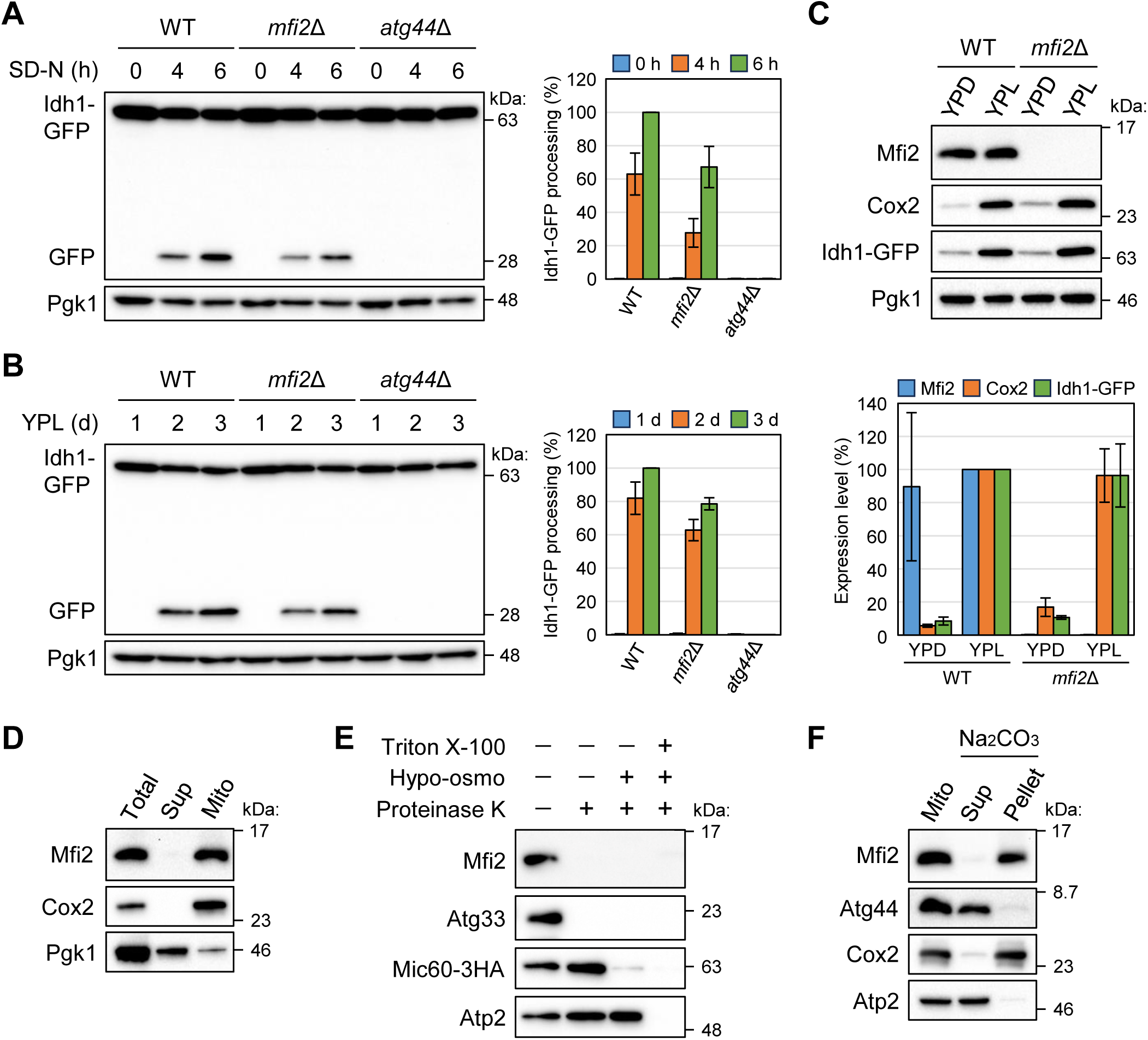
Mfi2 is a mitochondrial outer membrane-resident protein partially required for mitophagy. (A and B) The indicated cells were cultured in YPL until mid-log phase and shifted to synthetic minimal medium lacking nitrogen (SD-N) (A) or continuously cultured in YPL (B). Cells were collected at the indicated time points, and Idh1-GFP processing was monitored by immunoblotting. Pgk1 was detected as a loading control. The value of WT at 6-h (A) or 3-d (B) time point was set to 100%. The results represent the mean and SD of three experiments. (C) WT and *mfi2*Δ cells were cultured in YPD medium until early-log phase and in YPL until mid-log phase. Endogenous expression levels of Mfi2 were analyzed by immunoblotting. Cox2 and Idh1-GFP were detected as mitochondrial marker proteins. The value of WT (YPL) was set to 100%. The results represent the mean and SD of three experiments. (D) Subcellular fractionation was conducted using WT cells. The total cell homogenate was fractionated by centrifugation to obtain a mitochondria-enriched pellet and supernatant. Cox2 and Pgk1 were detected as markers of the mitochondria and cytosol, respectively. (E) Isolated mitochondria from cells expressing Mic60-3HA were treated with (+) or without (–) proteinase K under different conditions. Hypo-osmotic swelling resulted in rupture of the mitochondrial outer membrane, and the detergent Triton X-100 lysed mitochondria. Atg33, Mic60-3HA, and Atp2 were detected as markers of the mitochondrial outer membrane, intermembrane space, and matrix, respectively. (F) Isolated mitochondria were treated with sodium carbonate and separated into the soluble supernatant and membrane pellet by ultracentrifugation. Cox2 and Atp2 were detected as markers of integral and peripheral membrane proteins, respectively.

We found two similarities between Atg44 and Mco12. First, these proteins share a low but significant sequence homology (Figure S1C). Second, AlphaFold prediction^25^ revealed a structural similarity between the full length of Atg44 and the N-terminal two-thirds of Mco12 (Figure S1D). Notably, Mco12 possesses a C-terminal disordered region that is absent in Atg44 (Figure S1D), suggesting that Mco12 has distinct features from Atg44. CLIME analysis^26^ indicated that Mco12 is less conserved among fungal species than Atg44 (Figure S2). On the basis of its similarity to Atg44 and the functions described below, we renamed Mco12 as “*m*ito*fi*ssin *2*” (Mfi2).

### Mfi2 is a mitochondrial outer membrane protein

To characterize Mfi2, we first generated antibodies to detect endogenous Mfi2 (Figure 1C). We found that unlike the typical mitochondrial proteins Cox2 and Idh1, Mfi2 protein levels did not differ significantly in fermentation (YPD) and respiration (YPL) media (Figure 1C). Next, we examined the subcellular localization of Mfi2 by biochemical fractionation. As expected, Mfi2 was detected in the mitochondria-enriched fraction along with Cox2, but not in the cytosol-enriched fraction containing Pgk1 (Figure 1D). We then analyzed the intramitochondrial localization of Mfi2 using a proteinase K (ProK) protection assay. Degradation of Mic60 (intermembrane space, IMS) and Atp2 (mitochondrial matrix) required ProK treatment in combination with hypo-osmotic swelling and/or Triton X-100 lysis, whereas ProK treatment alone caused degradation of Mfi2 and the outer membrane protein Atg33 (Figure 1E). We also treated the mitochondrial fraction with sodium carbonate and separated membrane and supernatant fractions by ultracentrifugation. Mfi2 was detected in the pellet fraction along with the integral membrane protein Cox2, but not in the supernatant fraction containing the IMS protein Atg44 and the peripheral membrane protein Atp2 (Figure 1F). This result suggests that Mfi2 is strongly associated with mitochondrial membrane. Taken together, we conclude that Mfi2 is a mitochondrial outer membrane-resident protein.

### Mfi2 possesses mitochondrial fission activity *in vivo*

Next, we analyzed the significance of the N-terminal Atg44-like and C-terminal disordered regions of Mfi2. We expressed GFP-fused full-length, C- and N-terminal truncated Mfi2 proteins (Figure 2A) and verified their expression (Figure 2B). As shown in Figure 2C, GFP-Mfi2 and the C-terminal truncated GFP-Mfi2(N66) were co-localized with the mitochondrial marker Om14-RFP, whereas the N-terminal truncated GFP-Mfi2(C33) was observed throughout the cytosol. These results indicate that the Atg44-like region, but not the disordered region, is required for mitochondrial localization of Mfi2.

**Figure 2.**
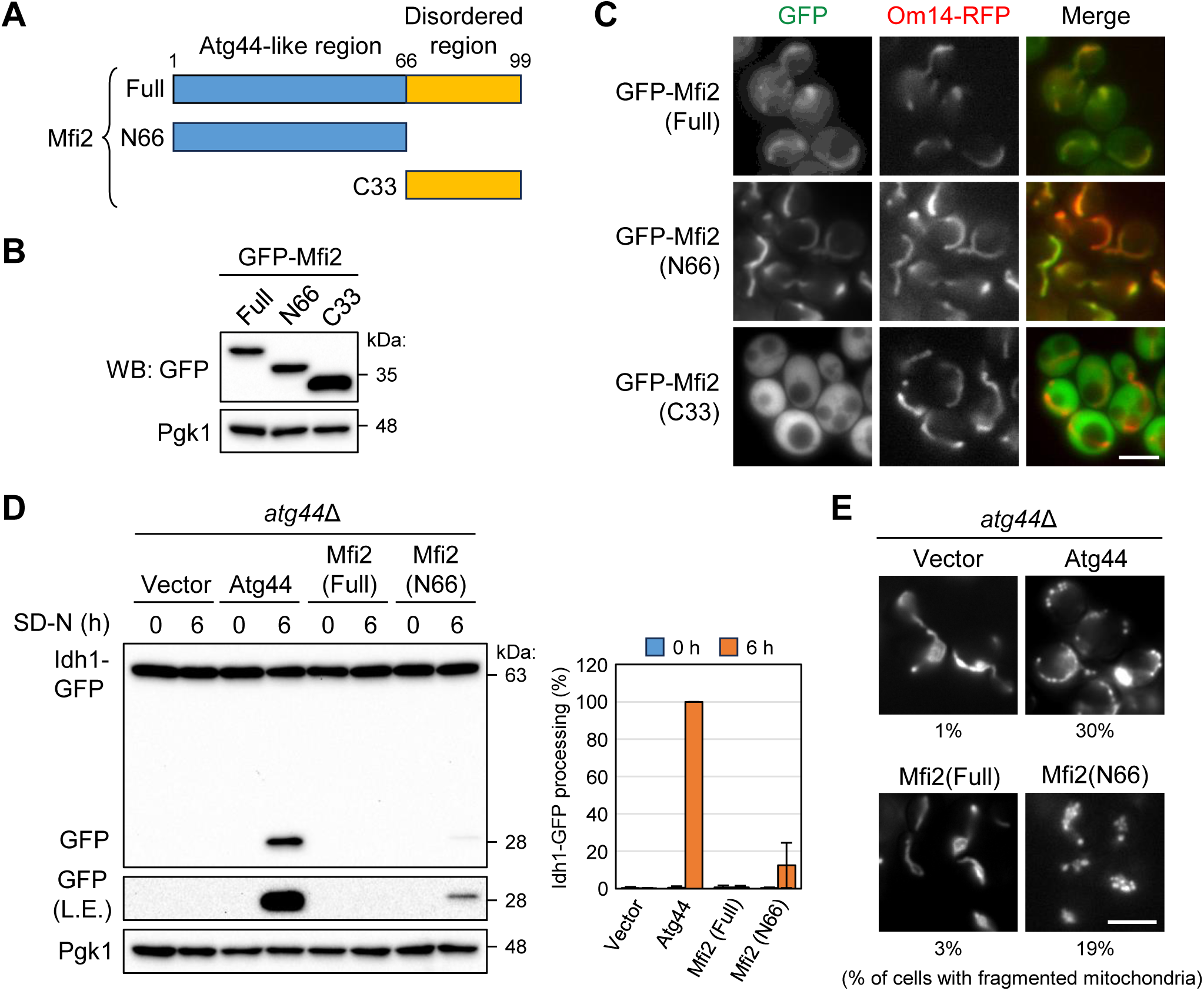
Mfi2 possesses mitochondrial fission activity *in vivo*. (A) Schematic representation of the N-terminal Atg44-like region (blue) and the C-terminal disordered region (orange) of Mfi2, and truncated forms. (B) Cells expressing GFP-fused Mfi2 derivatives were cultured in YPD until early-log phase, and their protein expression was verified by immunoblotting. (C) Cells expressing GFP-fused Mfi2 derivatives were cultured in YPD until early-log phase and their localization was analyzed by fluorescence microscopy. Om14-RFP was detected as mitochondrial marker. Scale bars, 4 μm. (D) *atg44*Δ cells expressing the indicated proteins were cultured in SML until mid-log phase and shifted to SD-N. Cells were collected at the indicated time points, and Idh1-GFP processing was monitored by immunoblotting. The value of Atg44 (6 h) was set to 100%. The results represent the mean and SD of three experiments. (E) WT cells and *atg44*Δ cells expressing the indicated proteins were cultured in SMD until mid-log phase and their mitochondrial morphology was analyzed by fluorescence microscopy. The percentage of cells with fragmented mitochondria is shown. Vector, n = 405; Atg44, n = 347; Mfi2(Full), n = 357; Mfi2(N66), n = 319. Scale bars, 4 μm.

We hypothesized that Mfi2(N66) functions similarly to Atg44, and therefore examined whether Mfi2(N66) can rescue the mitophagy defect of *atg44*Δ cells. Overexpression of full-length Mfi2 did not rescue mitophagy of *atg44*Δ cells, whereas overexpression of Mfi2(N66) slightly rescued the mitophagy defect (Figure 2D). In addition, overexpression of Mfi2(N66), but not full-length Mfi2, induced mitochondrial fragmentation in *atg44*Δ cells, which typically exhibit enlarged mitochondria^20–22^ (Figure 2E). These data demonstrate that Mfi2 possesses mitofissin activity *in vivo*.

### Mfi2 possesses lipid membrane-binding and -fission activity *in vitro*

To characterize Mfi2 *in vitro*, we prepared recombinant maltose-binding protein (MBP)-fused Mfi2 and MBP-Atg44 as a control (Figure S3A). Size-exclusion chromatography coupled with multi-angle light scattering (SEC-MALS) analysis estimated the molar masses of soluble MBP-Atg44 and MBP-Mfi2 as 1.16 and 0.99 MDa, respectively (Figure S3B). Their calculated molecular masses are 50 and 54 kDa, respectively, indicating that these recombinant proteins are highly oligomerized in solution. Using the recombinant proteins, we first examined whether Mfi2 binds to lipid membranes. Both Alexa Fluor 488 (AF488)-labeled MBP-Atg44 and AF647-MBP-Mfi2 accumulated on a giant unilamellar vesicle (GUV) derived from a lipid film with a phospholipid composition similar to that of the inner mitochondrial membrane (Figure 3A), indicating that Mfi2 also possesses lipid membrane-binding activity. Importantly, the membrane binding of both Mfi2 and Atg44 was severely impaired when cardiolipin (CL) was reduced or absent in the phospholipids, suggesting that their membrane binding *in vitro* requires CL (Figures 3A and S3C). We then prepared lipid nanotubes fluorescently labeled with liss Rhod PE^20,27^ and used them to analyze the membrane fission activity of Mfi2. Upon application of AF647-MBP-Mfi2 as well as AF488-MBP-Atg44, most of the nanotubes underwent fission at multiple sites. This fission event was strongly dependent on the presence of a high concentration of CL (Figure 3B). These data indicate that Mfi2 possesses membrane fission activity comparable to that of Atg44.

**Figure 3.**
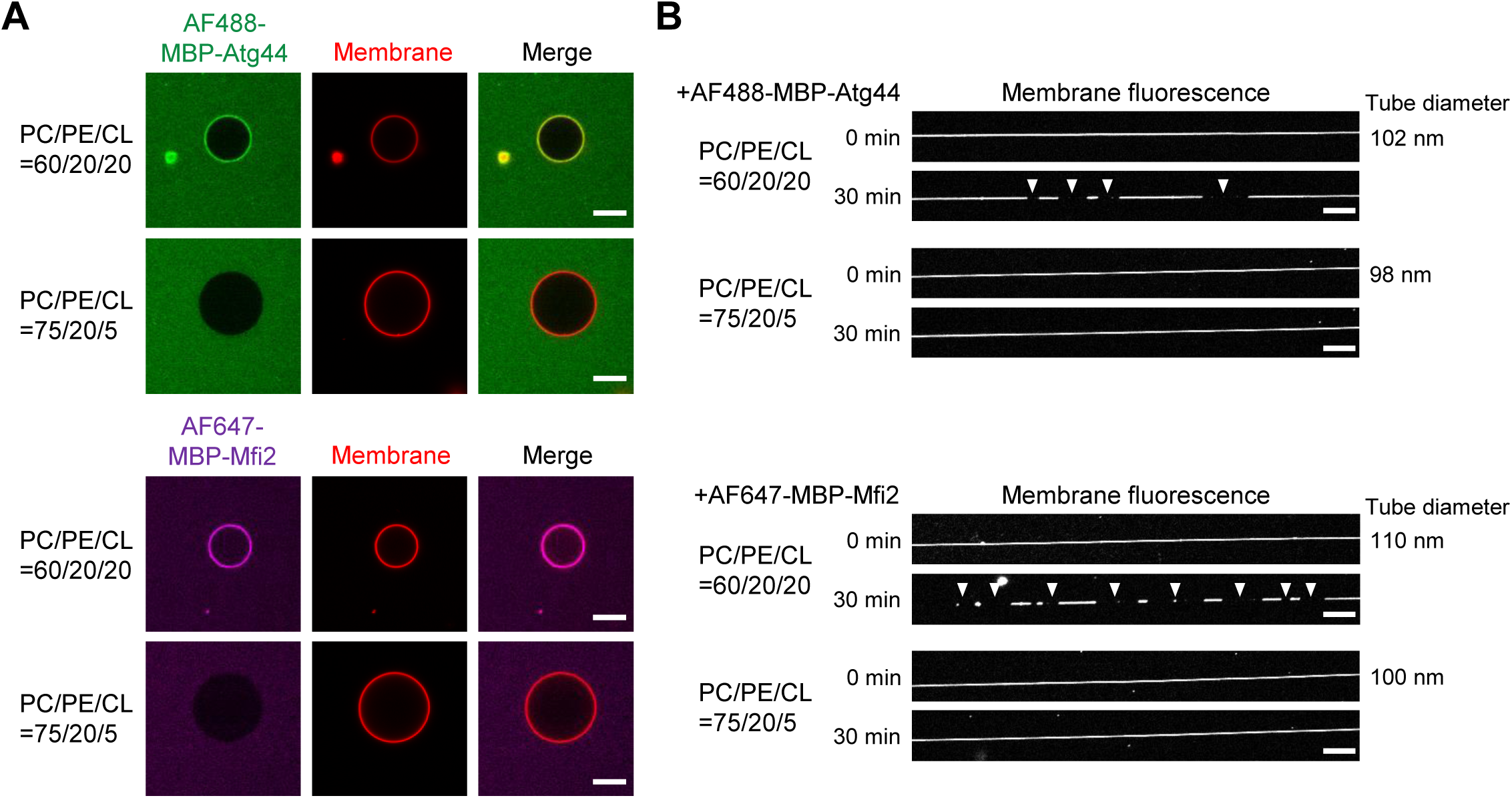
Mfi2 possesses membrane-binding and -fission activity *in vitro*. (A) Membrane binding of MBP-Atg44 and MBP-Mfi2. Membrane binding was examined by confocal laser scanning microscopy using fluorescently-labeled proteins and GUVs labeled with liss Rhod PE. Scale bars, 10 μm. (B) Membrane fission by MBP-Atg44 and MBP-Mfi2. Fission of lipid nanotubes was observed by confocal laser scanning microscopy using fluorescently-labeled proteins and lipid nanotubes labeled with liss Rhod PE. Positions of fission are marked with arrowheads. Scale bars, 10 μm.

### Mfi2 and Dnm1 independently contribute to mitophagy-associated mitochondrial fission

Dnm1 is the first identified mitochondrial fission factor,^28,29^ but it is dispensable for mitophagy.^17,18,20^ Mfi2 is also not essential for mitophagy (Figures 1A and 1B). These features led us to hypothesize that Mfi2 and Dnm1 redundantly contribute to mitochondrial fission during mitophagy. Therefore, we examined mitophagy in *mfi2*Δ *dnm1*Δ double mutant cells. As shown in Figure 4A, *mfi2*Δ and *dnm1*Δ single mutant cells exhibited slightly decreased mitophagy compared with WT cells, whereas the *mfi2*Δ *dnm1*Δ double mutant cells displayed a marked decrease in mitophagy. These results suggest that Mfi2 and Dnm1 independently contribute to mitophagy.

**Figure 4.**
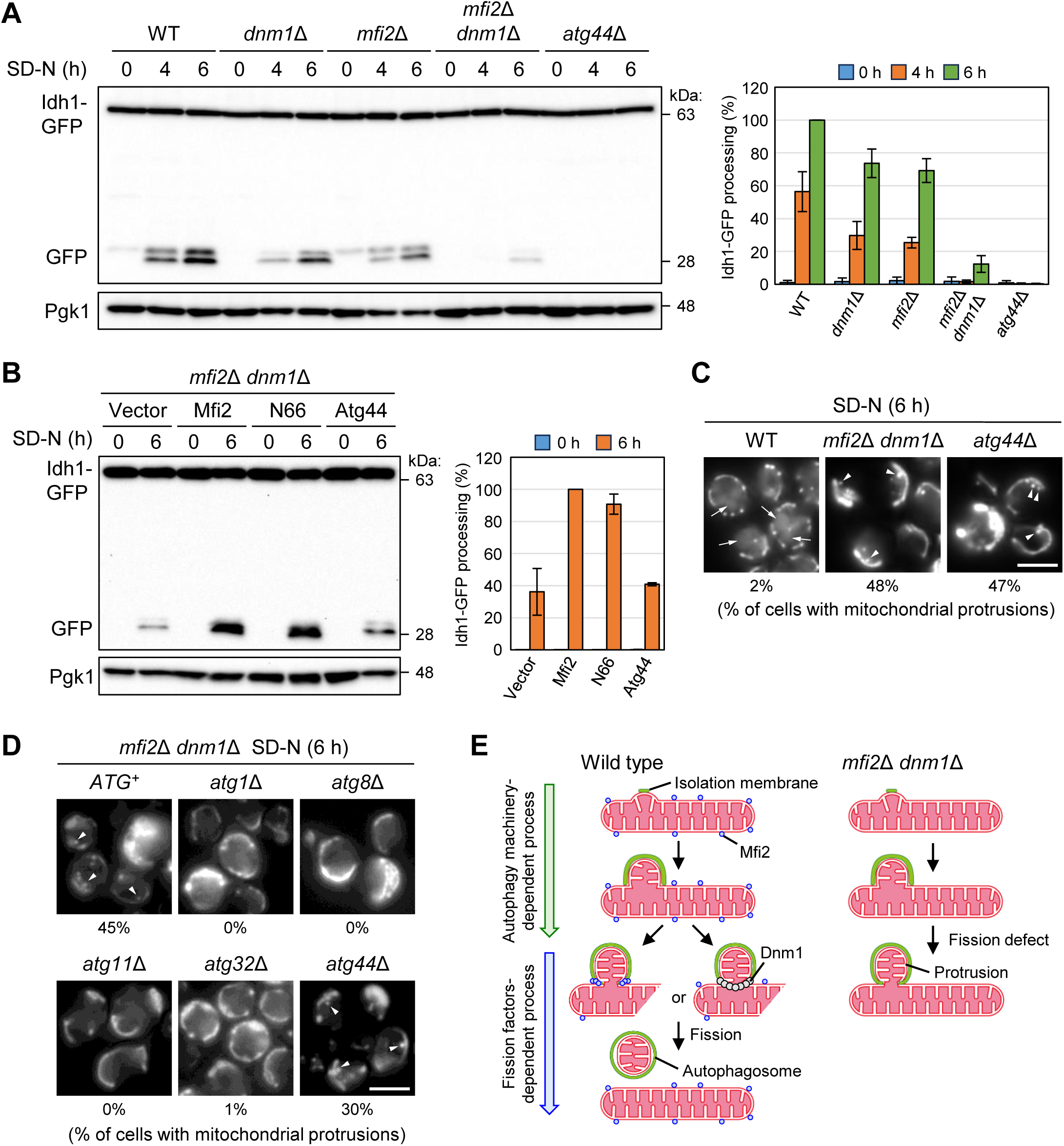
Mfi2 and Dnm1 independently contribute to mitophagy-associated mitochondrial fission. (A and B) The indicated cells were cultured in YPL (A) or SML (B) until mid-log phase and shifted to SD-N. Cells were collected at the indicated time points, and Idh1-GFP processing was monitored by immunoblotting. The value of WT at 6-h (A) or *dnm1*Δ *mfi2*Δ cells expressing Mfi2 at 6-h (B) time point was set to 100%. The results represent the mean and SD of three experiments. (C and D) The indicated cells were cultured in YPL until mid-log phase and shifted to SD-N for 6 h. The absence or presence of mitochondrial protrusions (arrowheads) was analyzed by fluorescence microscopy. Vacuolar GFP accumulation is indicated by arrows (WT in C). The percentage of cells with mitochondrial protrusions was quantified. WT, n = 258; *dnm1*Δ *mfi2*Δ, n = 211; *atg44*Δ, n = 251 in (C). *ATG*^+^, n = 331; *atg1*Δ, n = 420; *atg8*Δ, n = 300; *atg11*Δ, n = 403; *atg32*Δ, n = 326; *atg44*Δ, n = 303 in (D). Scale bars, 4 μm. (E) Schematic model of mitochondrial fission upon induction of mitophagy. See the main text for details.

Taking advantage of the impaired mitophagy in *mfi2*Δ *dnm1*Δ cells, we examined two Mfi2 orthologs from *Ashbya gossypii* (Ag) and *Debaryomyces hansenii* (Dh) (Figures S2 and S4A). Despite structural differences in the C-terminal regions predicted by AlphaFold (Figure S4B), expression of AgMfi2 or DhMfi2 restored mitophagy in *mfi2*Δ *dnm1*Δ cells to levels comparable to that of Mfi2. These results suggest that the C-terminal region of Mfi2 does not play a primary role in its function in mitophagy. We also compared the functions of Mfi2(N66) and Atg44 with Mfi2. Expression of Mfi2 or Mfi2(N66) restored mitophagy in *mfi2*Δ *dnm1*Δ cells at the same level, whereas expression of Atg44 did not (Figure 4B). These results suggest that the C-terminal region of Mfi2 is dispensable for its function and that Atg44, despite comparable *in vitro* activity, cannot fulfill the role of Mfi2 acting from outside the mitochondria. Taken together with the result that full-length Mfi2 cannot restore mitophagy in *atg44*Δ cells (Figure 2D), we conclude that both inner (Atg44) and outer (Mfi2) mitofissins are required for efficient mitophagy.

Next, we investigated why mitophagy is impaired in *mfi2*Δ *dnm1*Δ cells. In WT cells, mitochondrial Idh1-GFP signals were accumulated in vacuoles upon induction of mitophagy (Figure 4C, arrows). By contrast, *mfi2*Δ *dnm1*Δ cells exhibited mitochondrial protrusions, which is a typical morphological phenotype of the mitochondrial fission defect during mitophagy observed in *atg44*Δ cells^20^ (Figure 4C, arrowheads). As in the case of the protrusions observed in *atg44*Δ cells, this protrusion formation in *mfi2*Δ *dnm1*Δ cells was completely suppressed by disruption of autophagy/mitophagy factors such as Atg1 (a protein kinase of the Atg1 core complex), Atg8 (a ubiquitin-like protein required for autophagosome formation), Atg11 (an adaptor for selective autophagy), and Atg32 (a receptor for mitophagy) (Figure 4D), indicating that the mitochondrial protrusion formation depends on the mitophagy process. Notably, mitochondrial protrusions were also observed in *mfi2*Δ *dnm1*Δ *atg44*Δ triple mutant cells without additive effects (Figure 4D). These results suggest that the portion of the mitochondrial protrusion is separated from the main body of the mitochondrion during mitophagy by the coordinated action of the three fission factors, Mfi2, Dnm1, and Atg44. Taken together, we conclude that Mfi2 and Dnm1, as well as Atg44, are required for completion of mitochondrial fission during mitophagy.

## DISCUSSION

The identification of mitofissin/Atg44 as a mitochondrial fission factor essential for mitophagy raised the question of whether Atg44-mediated action from the IMS is sufficient for mitochondrial fission. In the present study, we identified Mfi2 as a mitochondrial outer membrane-resident mitofissin and found that simultaneous disruption of Mfi2 and Dnm1 resulted in a severe defect in mitophagy, suggesting that Mfi2 and Dnm1 independently contribute to mitochondrial fission during mitophagy. The mitochondrial fission defect in *mfi2*Δ *dnm1*Δ cells, even in the presence of Atg44, also suggests that Atg44 alone is not sufficient to complete mitochondrial fission. Thus, this study demonstrates that mitochondrial fission during mitophagy requires both “inner” and “outer” fission factors. The IMS mitofissin Atg44 promotes mitophagy-associated fission in coordination with the outer membrane protein mitofissin Mfi2 or the dynamin-related protein Dnm1, both acting from outside the mitochondria.

On the basis of our results and previous reports, we propose a model for mitochondrial fission during mitophagy in yeast (Figure 4E). Upon induction of mitophagy, isolation membranes emerge on mitochondria and extend along the mitochondrial surface.^18^ This process depends on the mitochondrial loading of the autophagy machinery, which is mediated by the interaction between the mitophagy receptor Atg32 and the adaptor Atg11.^20,30^ The extension of isolation membranes leads to the formation of a mitochondrial bud by an as yet unidentified mechanism, apparently independently of known mitochondrial fission factors (Figure 4D). Atg44 binds to mitochondrial membranes from the inside,^20,22^ while Mfi2 and Dnm1 bind to mitochondrial membranes from the outside. The coordinated actions of Atg44 with either Mfi2 or Dnm1 ultimately lead to mitochondrial fission, although the exact timing of this fission event remains to be determined. Finally, the mitochondrial fragment generated by this fission event is engulfed within the autophagosome. In the absence of either the internal action of Atg44 or the external action of Mfi2/Dnm1, the budded portion remains connected to the main body of the mitochondrion, resulting in protrusion formation.

Although this study advances our understanding of mitochondrial fission during mitophagy, several important questions remain unanswered. First, it is unclear how mitochondrial fission is facilitated in organisms that possess Atg44 orthologs but lack Mfi2 orthologs (e.g., *Schizosaccharomyces pombe* and most filamentous fungi) (Figure S2). In such organisms, it is possible that Atg44 orthologs can mediate both inner and outer mitochondrial membrane fission and/or that Dnm1 orthologs have sufficient activity for outer mitochondrial membrane fission. Comparative analysis of these orthologs across different species may provide insights that answer this question. Second, the importance of the C-terminal region of Mfi2 is unclear. The observation that the C-terminally truncated form of Mfi2 [Mfi2(N66)] is able to partially act like the IMS mitofissin Atg44 (Figures 2D and 2E) suggests that the presence of the C-terminal region prevents Mfi2 import into mitochondria. However, the majority of Mfi2(N66) was localized to the mitochondrial outer membrane, and even if present at very low levels in the IMS, it could not be detected (our unpublished data). Interestingly, Mfi2 orthologs such as AgMfi2 and DhMfi2 have different C-terminal structures, but they can rescue the function of Mfi2 (Figure S4C). This result suggests that the C-terminal region of Mfi2 has an unidentified but important role, although its structure has varied during evolution. One possible role of the C-terminal region of Mfi2 is to make Mfi1 localize strictly to the outer membrane. Third, it remains unclear why the *in vitro* lipid membrane binding of Mfi2 depends on a high concentration of CL in the lipid composition. This result seems counterintuitive given the observed localization of Mfi2 to the mitochondrial outer membrane and the fact that CL is predominantly contained in the mitochondrial inner membrane (∼20% in the inner membrane and ∼3% in the outer membrane).^31^ However, previous reports have shown that CL externalization from the inner to the outer membrane serves as a mitophagy signal.^32,33^ In addition, Drp1 has been shown to bind to CL, and this interaction is thought to be critical for its recruitment to mitochondria under stress conditions.^34^ These findings raise the possibility that Mfi2 functions in response to CL accumulation at the outer membrane of mitochondrial fission sites. Future studies are needed to test this hypothesis. Fourth, it is not yet clear whether Mfi2 and Dnm1 function cooperatively or independently for mitochondrial fission during mitophagy. In contrast to the enlarged mitochondrial morphology observed in *atg44*Δ and *dnm1*Δ cells,^20–22^ *mfi2*Δ cells display normal mitochondrial morphology under non-mitophagy-inducing conditions (Figure S1B). This observation suggests that Mfi2 and Dnm1 act independently, at least under homeostatic conditions, but further studies are needed to clarify this issue.

In summary, this study identifies Mfi2 as a mitochondrial outer membrane-type mitofissin and provides a compelling explanation for why Dnm1 is dispensable for mitophagy. Further studies will advance our understanding of how mitophagy-associated mitochondrial fission is orchestrated in yeast, and shed light on mitochondrial fission in higher eukaryotes where mitofissin counterparts have not yet been identified, although they are likely to exist. Furthermore, the discovery of mitofissin-like proteins required for the fission of other organelles, such as the endoplasmic reticulum and peroxisomes, would be of great interest, and our findings may provide a useful framework for such studies.

## STAR⍰METHODS

### RESOURCE AVAILABILITY

#### Lead contact

Further inquiries and requests for materials should be directed to the lead contact, Tomotake Kanki.

#### Materials availability

All requests for resources and reagents including plasmids and cell lines are available from the lead contact subject to a Materials Transfer Agreement.

### EXPERIMENTAL MODEL AND SUBJECT DETAILS

#### Yeast strains

The *Saccharomyces cerevisiae* strains used in this study are listed in Table S1. Gene deletion and tagging were performed as described previously.^35–37^

### METHODS DETAILS

#### Yeast culture conditions

Yeast cells were cultured at 30°C in rich medium (YPD: 1% yeast extract, 2% peptone, and 2% glucose), lactate medium (YPL: 1% yeast extract, 2% peptone, and 2% lactate), or synthetic minimal medium with glucose (SMD: 0.67% yeast nitrogen base, 2% glucose, and amino acids) or lactate (SML: 0.67% yeast nitrogen base, 2% lactate, and amino acids). Nitrogen starvation experiments were performed in synthetic minimal medium lacking nitrogen (SD-N: 0.17% yeast nitrogen base without amino acids and ammonium sulfate, and 2% glucose).

#### Fluorescence microscopy

Yeast cells expressing fluorescent proteins were cultured in the indicated media. Fluorescence images were captured using a Nikon Ti2 Eclipse microscope with a Plan Apo Lambda 100× oil objective lens and a CCD camera (MD-695, Molecular Devices) and analyzed using MetaMorph 7 (Molecular Devices).

#### Mitophagy assay

To monitor mitophagy, the Om45-GFP or Idh1-GFP processing assay was performed as previously described.^5,24^ In brief, cells were cultured in YPL or SML medium until mid-log phase, shifted to SD-N medium, and collected at the indicated time points. Cell lysates equivalent to OD_600_ = 0.2 units of cells were subjected to immunoblotting analysis.

#### Plasmids

To construct Mfi2 and Mfi2(N66) expression plasmids under the control of the *CUP1* or *TDH3* promoter, their coding regions were amplified by PCR and cloned into the BamHI-XhoI sites of pCu416^38^ or pTDH416.^20^ To construct AgMfi2 and DhMfi2 expression plasmids, their coding genes were artificially synthesized (Eurofins) and cloned into the BamHI-XhoI sites of pCu416. To construct a GST-fused truncated Mfi2 expression plasmid, the 26-99 a.a. coding region was amplified by PCR and cloned into the BamHI-XhoI sites of pGEX-4T-1 (Promega). Atg44 expression plasmids (pCu416-ATG44 and pTDH416-ATG44) were previously described.^20^ To construct Atg44 and Mfi2 expression plasmids for *in vitro* experiments, their coding genes were amplified by PCR and assembled into the downstream of the PreScission protease recognition sequence in a modified pET15b vector by using NEBuilder HiFi DNA assembly kit. A linker sequence (GlyGlyGlySerGlyGlyGlySer) was inserted between the maltose-binding protein (MBP) and PreScission protease recognition sequences. For fluorescent labeling, a cysteine residue was inserted into the linker sequence by PrimeSTAR Max mutagenesis (TaKaRa).

#### Antibodies

Anti-Mfi2 antibodies were produced by immunizing rabbits with recombinant GST-fused truncated Mfi2 (26-99 a.a.), followed by affinity purification with the same recombinant proteins transferred onto a polyvinylidene difluoride (PVDF) membrane (Merck Millipore). Anti-GFP (Takara Bio, 632380, RRID:AB_10013427), anti-Pgk1 (Thermo Fisher Scientific, 459250, RRID:AB_2532235), anti-Atp2 (Abcam, ab128743, RRID:AB_2810299), anti-HA (Sigma-Aldrich, H9658, RRID:AB_260092), anti-Cox2 (MitoScience, MS419, RRID:AB_1618187), anti-Atg33,^20^ and anti-Atg44^20^ were used for immunoblotting.

#### Immunoblotting analysis

Protein samples from yeast cells were resuspended in sodium dodecyl sulfate (SDS) sample buffer (50 mM Tris-HCl, pH 6.8, 10% glycerol, 2% SDS, 5% 2-mercaptoethanol, and 0.1% bromophenol blue), incubated at 42°C for 1 h, and subjected to SDS-polyacrylamide gel electrophoresis (PAGE). Proteins were transferred from polyacrylamide gels to PVDF membranes using transfer buffer (25 mM Tris, pH 8.3, 192 mM glycine, 20% methanol). The membranes were blocked with phosphate-buffered saline (PBS) with Tween-20 (PBS-T; 10 mM PO_4_^3-^, pH 7.4, 140 mM NaCl, 2.7 mM KCl, and 0.05% Tween-20) containing 5% skim milk for 1 h. The membranes were incubated with primary antibodies in PBS-T containing 2% skim milk overnight at 4°C and washed three times with PBS-T. The membranes were then incubated with secondary antibodies (Peroxidase-conjugated AffiniPure Goat Anti-Rabbit IgG or Peroxidase-conjugated AffiniPure Goat Anti-Mouse IgG) in PBS-T containing 2% skim milk for 1 h at room temperature and washed three times with PBS-T. Chemiluminescence signals were detected using ChemiDoc XRS+ (Bio-Rad) and analyzed using Image Lab software (Bio-Rad).

#### Isolation of mitochondria, sodium carbonate treatment, and ProK protection assay

Yeast cells were cultured in YPL until mid-log phase and resuspended in DTT buffer (10 mM dithiothreitol, 0.1 M Tris-HCl, pH 9.3) for 30 min at 30°C. Cells were collected and converted to spheroplasts in sorbitol buffer (1.2 M sorbitol, 20 mM KH_2_PO_4_, pH 7.4) with Zymolyase 100T (Nacalai tesque). Spheroplasts were collected by centrifugation (1000 × g for 10 min at 4°C) and resuspended in ice-cold homogenization buffer (0.6 M sorbitol, 20 mM HEPES, pH 7.4) and homogenized in a Potter-Elvehjem homogenizer. The cell homogenate was centrifuged at 1000 × g for 10 min at 4°C. The supernatant was centrifuged at 6500 × g for 10 min at 4°C and the pellet was collected as the mitochondrial fraction. Isolated mitochondria were separately suspended in ice-cold 0.1 M sodium carbonate (pH 11.0), incubated for 30 min on ice, and then centrifuged at 100,000 × g for 30 min at 4°C. Proteins in the pellet and supernatant fractions were precipitated by adding 10% TCA. Isolated mitochondria were separately suspended in ice-cold homogenization buffer, hypotonic buffer (20 mM HEPES, pH 7.4), or hypotonic buffer with 0.5% Triton X-100 and treated with ProK (200 μg/ml) for 30 min on ice. The ProK reaction was stopped by adding 10% trichloroacetic acid (TCA). TCA-precipitated proteins were washed with acetone and subjected to immunoblotting.

#### Preparation of recombinant Atg44 and Mfi2

Atg44 and Mfi2 were overexpressed as MBP-fusion proteins in *Escherichia coli* C41(DE3) cells (New England Biolabs). Cells were grown in Luria-Bertani medium and protein expression was induced with 0.5 mM isopropyl-β-D-thiogalactopyranoside (IPTG). After culture at 30°C for 3 hours, cells were harvested by centrifugation and suspended in P buffer (PBS supplemented with 1.0 M NaCl). The suspension was supplemented with 1.0 mM phenylmethylsulfonyl fluoride and lysed by sonication. The supernatant of the lysate after centrifugation was loaded onto Amylose resin (New England Biolabs) pre-equilibrated with P buffer. The resin was washed extensively with P buffer and equilibrated with PBS buffer. The bound proteins were eluted with PBS buffer supplemented with 10 mM maltose, concentrated with centrifugal device, Amicon Ultra-15 (Merck Millipore) and then subjected to SEC using Superose 6 Increase 10/300 GL column (Cytiva) in HEPES buffer (20 mM HEPES-NaOH, pH 7.0 and 150 mM NaCl). The eluates were concentrated with a centrifugal device and frozen at −80[for storage.

#### SEC-MALS

SEC-MALS was conducted with a chromatography system connected in-line to a DAWN HELEOS II (Wyatt) for light scattering and RI-501 (Shodex) for differential refractive index measurements. 100 μl of 50 μM MBP-Atg44 or MBP-Mfi2 was subjected to SEC using Superose 6 Increase 10/300 GL column in HEPES buffer at room temperature. Molar masses of elution peaks were calculated with Astra software (Wyatt) using the values of light scattering, differential refractive index and ultraviolet absorption at 280 nm.

#### Fluorescent labeling

Cys-introduced MBP-Atg44 was mixed with Alexa Fluor 488 C_5_-maleimide (Invitrogen) dissolved in dimethyl sulfoxide (DMSO) at an equivalent molar ratio. Cys-introduced MBP-Mfi2 was mixed with Alexa Fluor 647 C_2_-maleimide (Invitrogen) dissolved in DMSO at an equivalent molar ratio. The mixture was incubated for 30 minutes at room temperature and dialyzed against HEPES buffer using mini dialysis kit (Cytiva) at 4[overnight to remove remaining fluorescent dyes.

#### Membrane binding experiments

1-palmitoyl-2-oleoyl-sn-glycero-3-phosphocholine (PC), 1-palmitoyl-2-oleoyl-sn-glycero-3-phosphoethanolamine (PE), L-a-phosphatidylinositol (PI), 1-palmitoyl-2-oleoyl-sn-glycero-3-phospho-L-serine (PS), 1-palmitoyl-2-oleoyl-sn-glycero-3-phosphate (PA), 1’,3’-bis[1,2-dioleoyl-sn-glycero-3-phospho]-glycerol (CL) and 1,2-dioleoyl-sn-glycero-3-phosphoethanolamine-N-(lissamine rhodamine B sulfonyl) (liss Rhod PE) were purchased from Avanti Polar lipids. GUVs were prepared by a gel-assisted swelling method as described elsewhere.^39^ Briefly, a thin layer of 5% poly(vinyl alcohol) (PVA) with an average molecular weight of 146,000-186,000 (Sigma-Aldrich) dissolved in water was applied on a coverslip pre-cleaned with water and ethanol and dried at 50[. 1.0 mM lipid solution in chloroform was applied to the PVA sheet and dried in a desiccator connected to a vacuum pump for 1 hour. The dried lipid film on the PVA sheet was then hydrated in 320 mM sucrose for at least 30 minutes at room temperature to produce GUVs. The solution and PVA sheet were transferred to a plastic tube and the PVA sheet was removed before the solution containing GUVs was used.

For membrane binding analysis, a small aliquot of the solution containing GUVs was added to HEPES buffer in glass bottom dish (MatTek) passivated with BSA (Wako). Fluorescently-labeled MBP-Atg44 or MBP-Mfi2 were then added to the solution at final concentrations of 2.0 μM and mixed gently. Confocal images were acquired at room temperature with a confocal laser scanning microscope, A1 LFOV (Nikon) with a 60× oil immersion objective lens (Nikon) using NIS-elements software (Nikon).

#### Tube fission experiments

Lipid nanotubes were prepared using excess membrane reservoirs as described previously.^20,27^ Fluorescently-labeled MBP-Atg44 or MBP-Mfi2 was applied to lipid nanotubes at final concentrations of 2.0 μM in HEPES buffer in 8 well chamber cover (Matsunami Glass) passivated moderately with BSA. Confocal images were acquired at room temperature with A1 LFOV confocal microscope with a 60× oil immersion objective lens using NIS-elements software. Tube diameter was estimated from fluorescence intensity as described previously.^20,27^

## ACKNOWLEDGMENTS

This study was supported in part by the Japan Society for the Promotion of Science grants JP23H04255 and JP24K01717 (to K.F.), JP24K09373 (to T.M.), JP23K20044, JP24H00060, JP25H00966, and JP25H01321 (to N.N.N.), JP19H05712, JP23K23878, JP24H02274, and JP25H01101 (to T.K.); AMED-CREST grant 23gm1710006s0101 (to T.K.); JST-CREST grant JPMJCR20E3 (to N.N.N.); the Takeda Science Foundation (to K.F.); Institute for Fermentation, Osaka (IFO) (to K.F.); the Noda Institute of Scientific Research (to K.F.). We thank Edanz (https://jp.edanz.com/ac) for editing a draft of this manuscript.

## AUTHOR CONTRIBUTIONS

K.F., T.M., N.N.N., and T.K. designed the experiments. K.F. performed experiments in yeast with the help of S.Y., K.I., T.F., and T.K. T.M. and N.N.N. performed *in vitro* experiments. All authors analyzed the data. K.F., T.M., T.F., N.N.N., and T.K. wrote the manuscript with contributions from all authors.

## Supplemental Figure Legends

**Figure S1.**
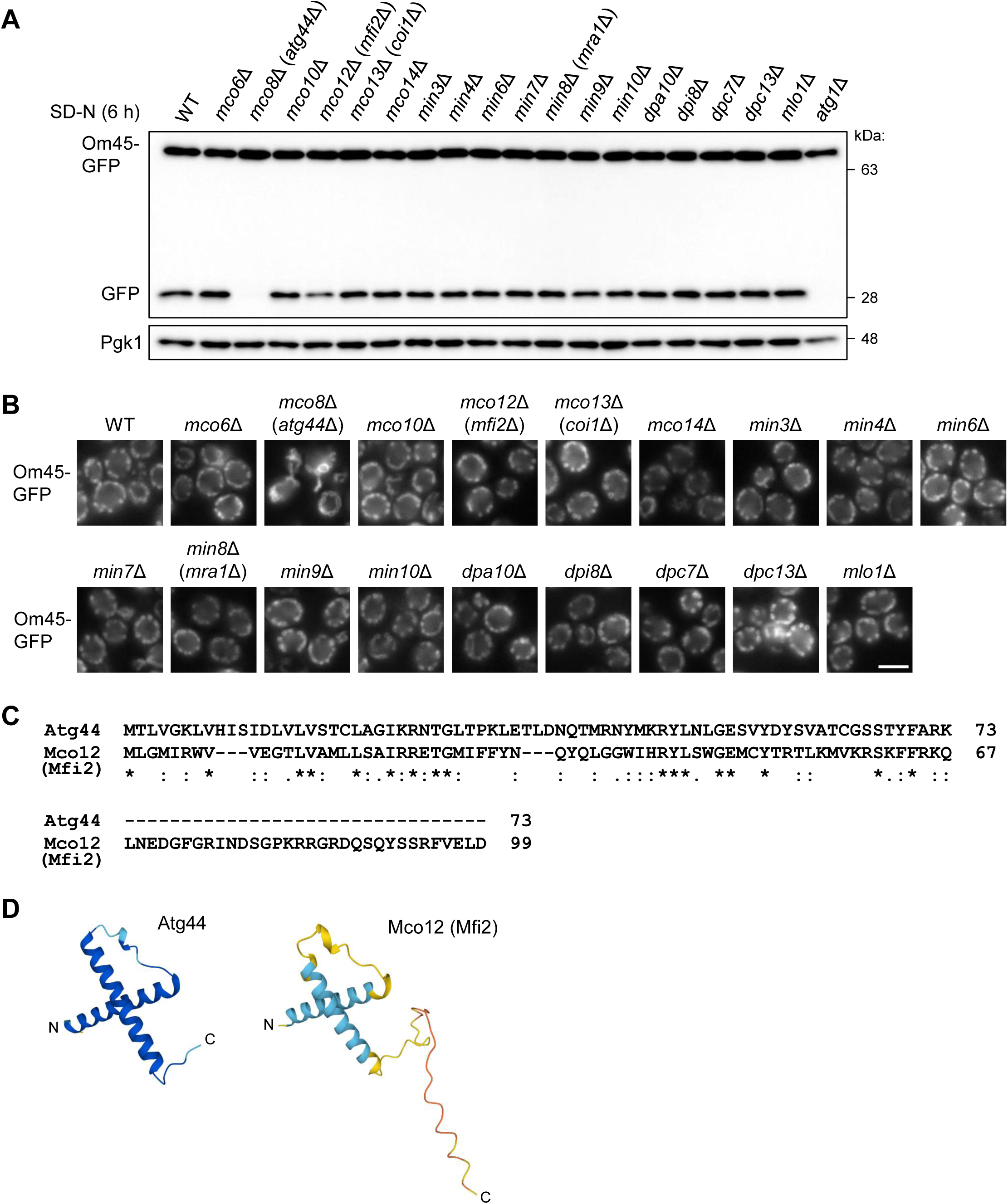
Identification of Mco12/Mfi2 as an Atg44-like protein, related to Figure 1. (A) The indicated cells were cultured in YPL until mid-log phase and shifted to SD-N. Cells were collected after 6 h, and Om45-GFP processing was monitored by immunoblotting. Pgk1 was detected as a loading control. (B) The indicated cells were cultured in YPL until mid-log phase, and their mitochondrial morphology was analyzed by fluorescence microscopy. More than 300 cells were analyzed, and representative images are shown. Scale bars, 4 μm. (C) Protein sequence alignment of Atg44 and Mco12/Mfi2. Asterisk, conserved residue; colon, residue with highly similar properties; period, residue with weakly similar properties. (D) AlphaFold-predicted structures of Atg44 and Mco12/Mfi2. N and C indicate the N- and C-termini, respectively.

**Figure S2.**
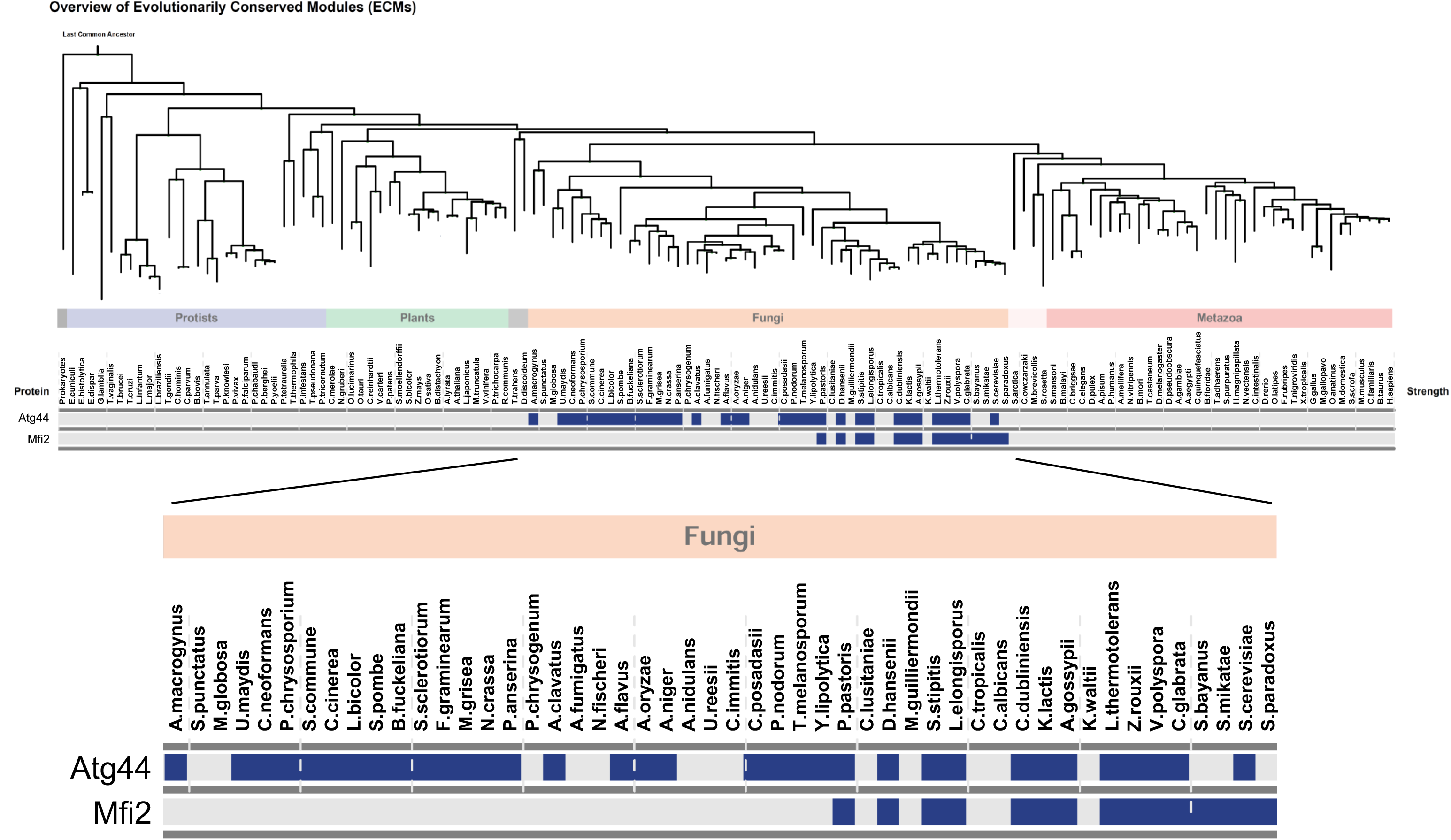
CLIME analysis of Atg44 and Mfi2, related to Figure 1. CLIME analysis (https://www.gene-clime.org/) shows less conservation of Mfi2 than of Atg44 among fungi.

**Figure S3.**
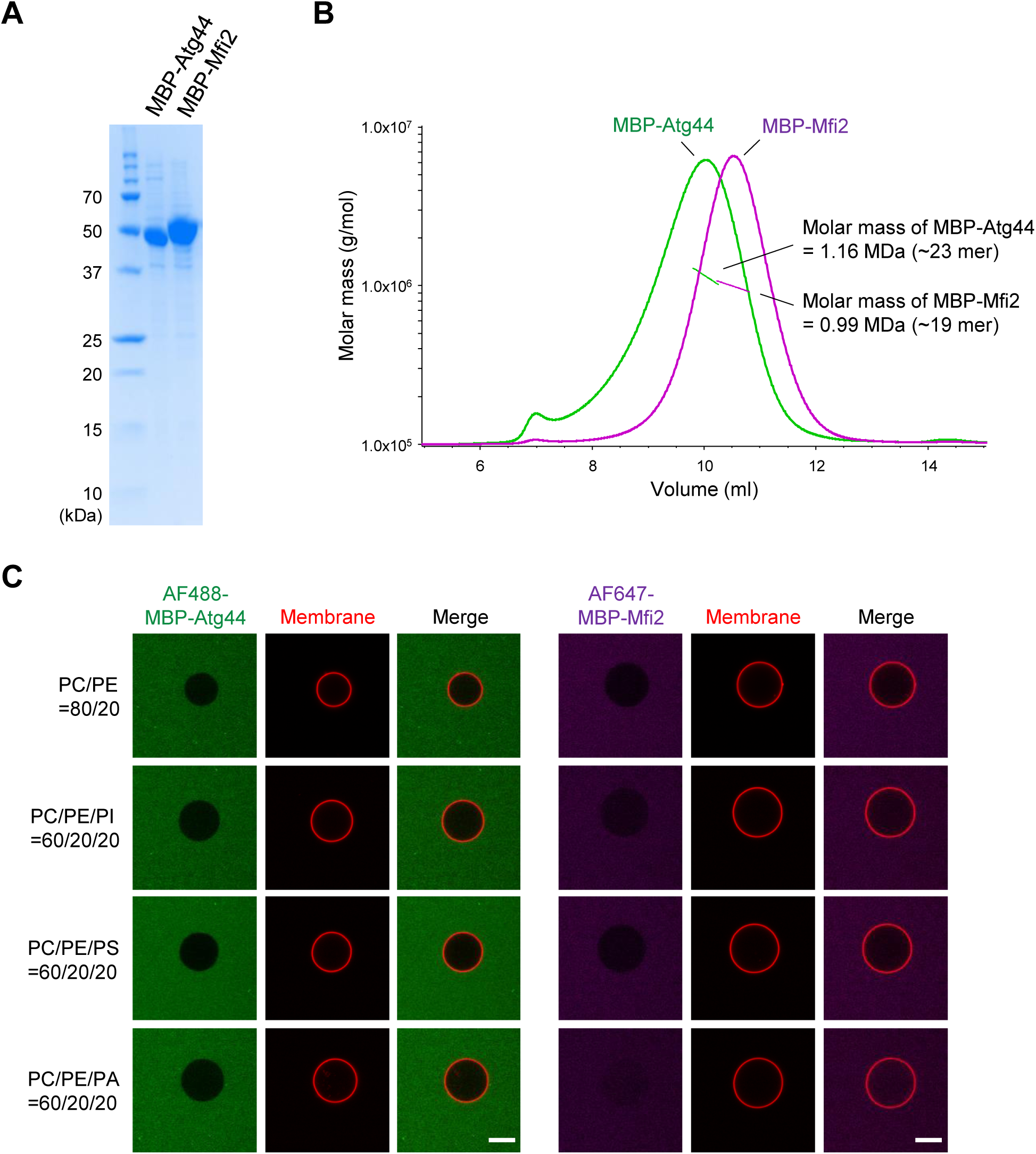
Sample preparation and membrane binding, related to Figure 3. (A) SDS-PAGE analysis of the purity of prepared samples. The gel was stained with Coomassie Brilliant Blue. (B) SEC-MALS analysis of the oligomerization states of MBP-Atg44 and MBP-Mfi2. Absorbance at 280 nm in SEC was shown as solid line. Estimated molar mass was shown as dot. (C) Membrane binding of MBP-Atg44 and MBP-Mfi2. Membrane binding was examined by confocal laser scanning microscopy using fluorescently-labeled proteins and GUVs labeled with liss Rhod PE. Scale bars, 10 μm.

**Figure S4.**
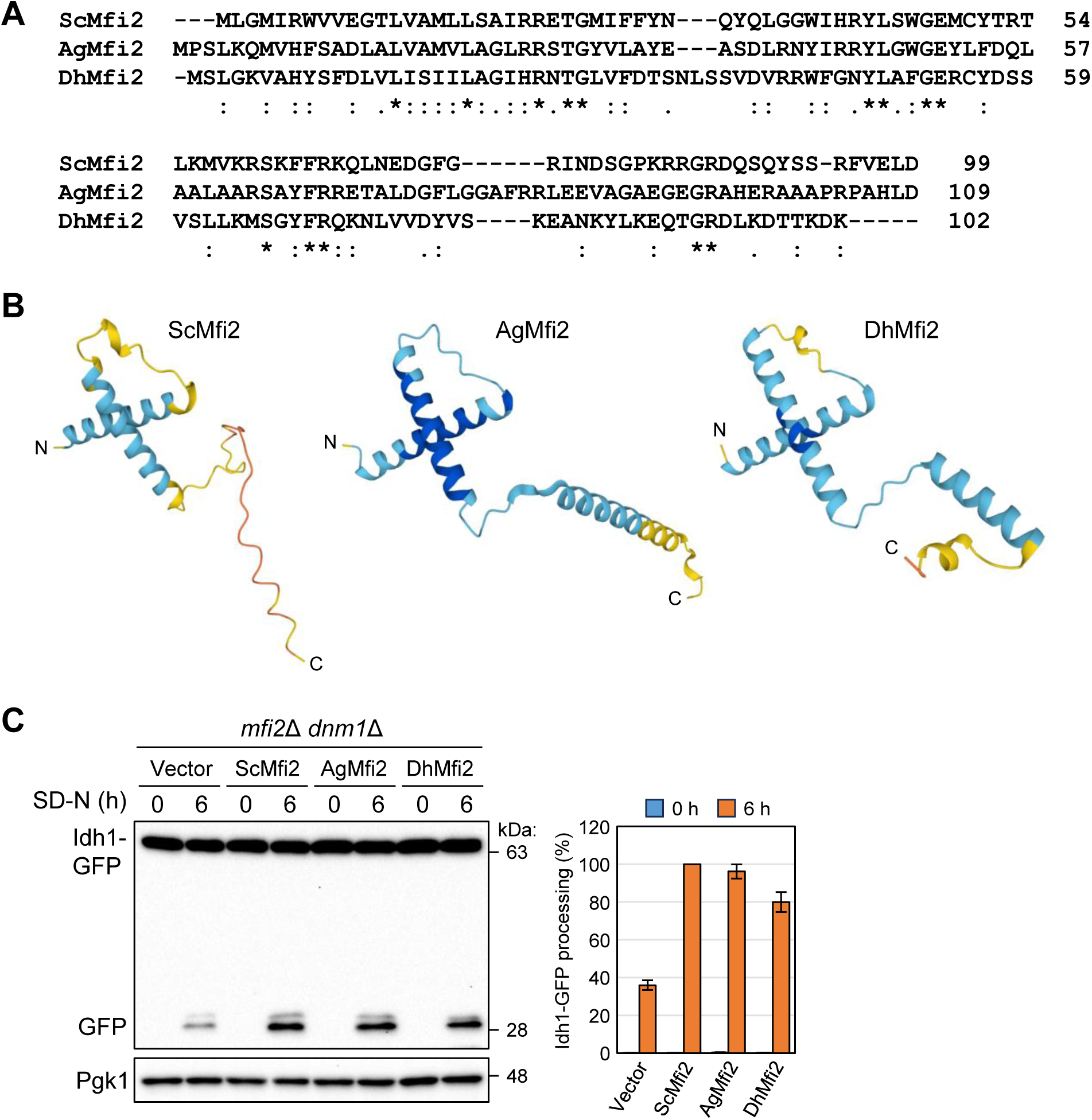
Exogenous expression of AgMfi2 or DhMfi2 rescues the impaired mitophagy in *dnm1*Δ *mfi2*Δ cells, related to Figure 4. (A) Protein sequence alignment of Mfi2 homologs. Sc, *Saccharomyces cerevisiae*; Ag, *Ashbya gossypii*; Dh, *Debaryomyces hansenii*. Asterisk, conserved residue; colon, residues with highly similar properties; period, residues with weakly similar properties. (B) AlphaFold2-predicted structures of Mfi2 homologs. (C) The *dnm1*Δ *mfi2*Δ cells expressing the indicated proteins were cultured in SML until mid-log phase and shifted to SD-N. Cells were collected at the indicated time points, and Idh1-GFP processing was monitored by immunoblotting. The value of ScMfi2 (6 h) was set to 100%. The results represent the mean and SD of three experiments.

**Table S1.**
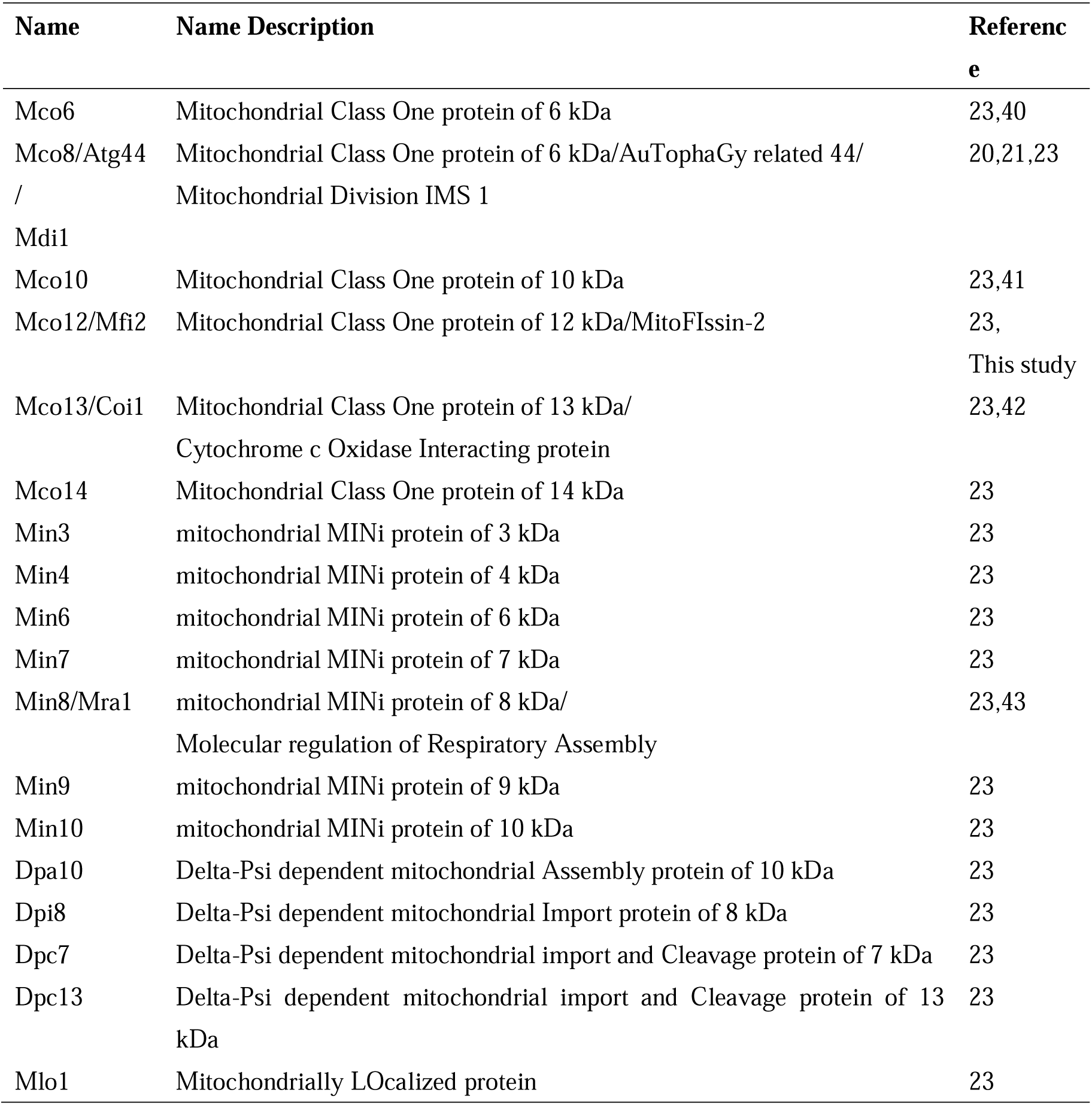
Mitochondrial small proteins analyzed in this study, related to Figure S1.

**Table S2.** Yeast strains used in this study, related to STARfllMETHODS.

| **Strain** | **Genotype** | **Reference** |
| --- | --- | --- |
| YKF27 | *IDH1-GFP::TRP1* | 30 |
| YKF92 | *atg44::kanMX IDH1-GFP::TRP1* | 20 |
| YKF250 | *IDH1-GFP::hphNT* | 20 |
| YKF251 | *atg44::kanMX IDH1-GFP::hphNT* | 20 |
| YKF253 | *dnm1::kanMX IDH1-GFP::hphNT* | 20 |
| YKF298 | *mfi2::kanMX IDH1-GFP::TRP1* | This study |
| YKF305 | *mfi2::natNT IDH1-GFP::hphNT* | This study |
| YKF313 | *mfi2::natNT dnm1::kanMX IDH1-GFP::hphNT* | This study |
| YKF322 | *IDH1-GFP::TRP1 MIC60-3HA::HIS3* | This study |
| YKF327 | *OM14-RFP::TRP1 natNT::PADH1-GFP-MFI2* | This study |
| YKF330 | *OM14-RFP::TRP1 natNT::PADH1-GFP-MFI2(N66)::HIS3* | This study |
| YKF336 | *OM14-RFP::HIS3 natNT::PADH1-GFP-MFI2(C33)* | This study |
| YKF375 | *mfi2::HIS3 dnm1::kanMX atg1::natNT IDH1-GFP::hphNT* | This study |
| YKF376 | *mfi2::HIS3 dnm1::kanMX atg8::natNT IDH1-GFP::hphNT* | This study |
| YKF377 | *mfi2::HIS3 dnm1::kanMX atg11::natNT IDH1-GFP::hphNT* | This study |
| YKF378 | *mfi2::HIS3 dnm1::kanMX atg32::natNT IDH1-GFP::hphNT* | This study |
| YKF379 | *mfi2::HIS3 dnm1::kanMX atg44::LEU2 IDH1-GFP::hphNT* | This study |

